# Local ecological knowledge, catch characteristics and evidence of elasmobranch depletions in Western Ghana

**DOI:** 10.1101/2021.01.14.426682

**Authors:** Issah Seidu, Lawrence K. Brobbey, Emmanuel Danquah, Samuel K. Oppong, David van Beuningen, Nicholas K. Dulvy

**Author notes:** Corresponding author ✉ Issah Seidu, Department of Wildlife and Range Management, Kwame Nkrumah University of Science and Technology, PMB, Kumasi – Ghana.

## Abstract

Local Ecological Knowledge has the potential to improve fishery management by providing new data on the fishing efforts, behavior, and abundance trends of fish and other aquatic animals. Here, we relied on local knowledge of fishers to investigate ecological factors that affect elasmobranch fishers‟ operations and the changes in stock status of sharks and rays from 1980 to 2020 in five coastal communities in Ghana. Data were gathered from fishers using participant observation, interviews, focus group discussions, and participatory rural appraisal techniques. The results revealed fisher‟s understanding of six main ecological variables, which are mostly applied to enhance their fishing operations: season and weather conditions, lunar phase, bait type, presence of seabirds and fish movement, color of seawater, and sea current. These ecological features have been applied over the years to enhance fishing operations as well as maximize fisher catch. Fishers reported a profound decline in shark and ray catch from 1980 to 2020 and attributed the decline in size, number, and composition of their catch to overfishing and Illegal, Unreported and Unregulated (IUU) fishing operations. In general, most shark and ray species were abundant in 1980 but have been severely depleted as of 2020, with the exception of Blue Shark (*Prionace glauca)* and Devil rays (*Mobula* spp), which were reported to be common by the interviewed fishers. The first species depleted were the Thresher sharks (Alopiidae), Tiger Shark (*Galeocerdo cuvier*), Blackchin Guitarfish (*Glaucostegus cemiculus*), and Lemon Shark (*Negaprion brevirostris*), which were depleted early in the 2000s. The next depletions of Hammerhead sharks (Sphyrnidae), Bull Shark (*Carcharhinus leucas*), Sand Tiger Shark (*Carcharias taurus*), Stingrays (*Fontitrygon* spp), and Spineback Guitarfish (*Rhinobatos irvinei*) occurred in the 2010s. We found Local Ecological Knowledge of fishers to be surprisingly consistent with scholarly knowledge and call for their inclusion in research, decision-making and management interventions by biologists and policy makers.

## 1. Introduction

The data needed for managing artisanal fisheries exploitation, like population trends and catch efforts, have traditionally been acquired from scientific surveys and monitoring methods (Castillo-Géniz *et al.,* 1998; Lunn and Dearden, 2006). However, in developing countries with high biodiversity and mixed-species fisheries, these time consuming and relatively expensive methods may be difficult to implement (Johannes *et al.,* 2000; Anadon *et al.,* 2009). With these difficulties in gathering data, researchers have to depend on alternative knowledge from local fishers to understand biological and ecological processes in the artisanal fishery setting (Johannes *et al.,* 2000; Moore *et al.,* 2010). There has also been historical recognition that wildlife and fisheries management can no longer depend only on empirical scientific data (Mangel *et al.,* 1996; McCool and Guthrie 2001; Riley *et al.,* 2002). Therefore, the incorporation of the social dimension in developing relevant management protocols is widely advocated (McCleery *et al.,* 2006; Miller, 2009).

Local ecological knowledge (LEK) has been described as practical, behavior-oriented, structured, dynamic, and based on long-term empirical observations (Ruddle, 2000). The application of LEK in fisheries management has attracted public interest (Olsson and Folke, 2001; Berkes *et al.,* 2008). It has been extensively used in fishery studies to characterize gear types and fishing efforts (Lam and Sadovy de Mitcheson, 2011), estimate bycatch (Silver and Campbell, 2005), and provide biological data such as reproduction and fish diet (Grant and Berkes, 2007; Rasalato *et al.,* 2010). LEK has also been used to study the population status and trends of fishes (Taylor *et al.,* 2011), lobsters (Eddy *et al.,* 2010), fisheries dynamics (Ainsworth *et al.,* 2008), trends in ecological processes (Poizat and Baran, 1997), and to detect extinctions (Dulvy and Polunin, 2004). Further, Silvano and Valbo-Jorgensen (2008) hypothesize that LEK of fishers may have the potential to improve fishery management through the provision of new information about the ecology, behavior, and abundance trends of fishes and other aquatic animals.

In contrast to specialized industrial fleets that have finite fishing seasons and are mostly regulated by policies, artisanal fisheries are generally flexible, dynamic and operate almost all year round, with little or no management structures (Salas *et al.,* 2007; Moore, 2010). Artisanal fisheries contribute 60% of animal protein consumed in Ghana, with average per-capita consumption between 20–25 kg per annum (Nunoo *et al.,* 2014; National Fisheries Management Plan, 2015). Ghana‟s fishery sector provides direct and indirect employment to 10% of Ghanaians, which translates into 2.6 million people of the current population (Finegold *et al.,* 2010; Nunoo *et al.,* 2014). Ghana has a 550 km coastline with about 90 lagoons and associated wetlands (deGraft-Johnson *et al.,* 2010). The extensive coastline, lagoons, and rivers support many artisanal fisheries. Ghanaian artisanal fisheries range from subsistence to small-scale commercial fishing, part-time to full-time, sedentary to migrant fishers, non-advanced, and non-differentiated, to highly specialized and differentiated forms of fishing operations (Demuynck, 1994; Aheto *et al.,* 2012). According to census data, there are about 12,000 artisanal canoes with over 124,000 fishers operating in 300 landing sites along Ghana‟s coastline (Amador *et al.,* 2006; CRC, 2013). The fishing effort is likely to be underestimated, since most fishers and canoes are not registered and operate illegally and may not have been counted in the census. Artisanal fishery contributes 80% to the total national annual marine fish landings by volume (CRC, 2013).

There is growing awareness of declines of shark and ray populations and efforts are underway to provide relevant data on their status and develop strategies to halt and reverse their decline (Dulvy *et al.,* 2014; Bräutigam *et al.,* 2015). Nevertheless, few data exist regarding shark and ray catches and their status in West Africa (Fowler *et al.,* 2005). A recent conservation strategy highlighted the urgent need to improve the collection, reporting, and analysis of information on sharks and rays throughout most of their range in West Africa, including Ghana, to guide improved fisheries management (Bräutigam *et al.,* 2015; Dulvy *et al.,* 2017). However, the small sizes of artisanal boats or canoes make on-board observation of catches and record keeping logistically impossible, as there is no room to accommodate observers (Moore, 2010). With these situations, in-situ monitoring and estimates of vessel catch, effort, by-catch, and operations may not be applicable in artisanal fisheries and as such these fisheries are typically data-poor (Kelleher, 2005; Salas *et al.,* 2007; Moore, 2010). Additionally, discards at sea are largely unaccounted for leading to information related to dead removals of by-catch (dead discards plus declared landings) being largely unavailable from artisanal fishers (Bonfil, 1994). Accordingly, national and global reported statistics on catch stock and fishing capacity may not yield accurate scientific data (Moore *et al.,* 2010). As a result of these challenges, the current and historical status of sharks and rays in Ghana, including their catch characteristics, composition, threats, and operations of fishers targeting these species are completely unknown. In an effort to contribute to the literature on retrospective assessment of artisanal elasmobranch fisheries, this study used local knowledge of fishers in Western Ghana to ascertain ecological factors that affect elasmobranch fishers‟ fishing operations, and historical and current records of the status of sharks and rays in Ghana.

Here we tackle the following questions: (i) how do elasmobranch fishers learn, understand, and apply ecological features to enhance their fishing operations?, (ii) have the catch composition and sizes of target species changed over time?, and (iii) have fishers catch abundance of commercially important shark and ray species change since the 1980s?

## 2. Methodology

First, we describe the five coastal communities of the study sites. Second, we describe the data collection methods, and; third, the data analysis methods.

### 2.1 Study sites

This study was conducted in five coastal communities, which are among the hotspots of elasmobranch fisheries in the Western Region of Ghana (Figure 1, Table 1). These communities were selected based on three main reasons: (i) fishing is exclusive to artisanal fishers; (ii) sharks and/or rays form a significant component of the catch; and (iii) fishers were willing to cooperate with the researchers for both landing and interview data. The communities are located in West Coast (≈95 km) and extend from the Ghana-Côte d‟Ivoire border to the Ankobra River estuary (deGraft-Johnson *et al.,* 2010). Tuesdays are observed as fishing holidays in all five study communities. Fishermen in these communities used Tuesdays to mend their gears and equipment, resolve conflicts, and carry out other social activities.

**Figure 1:**
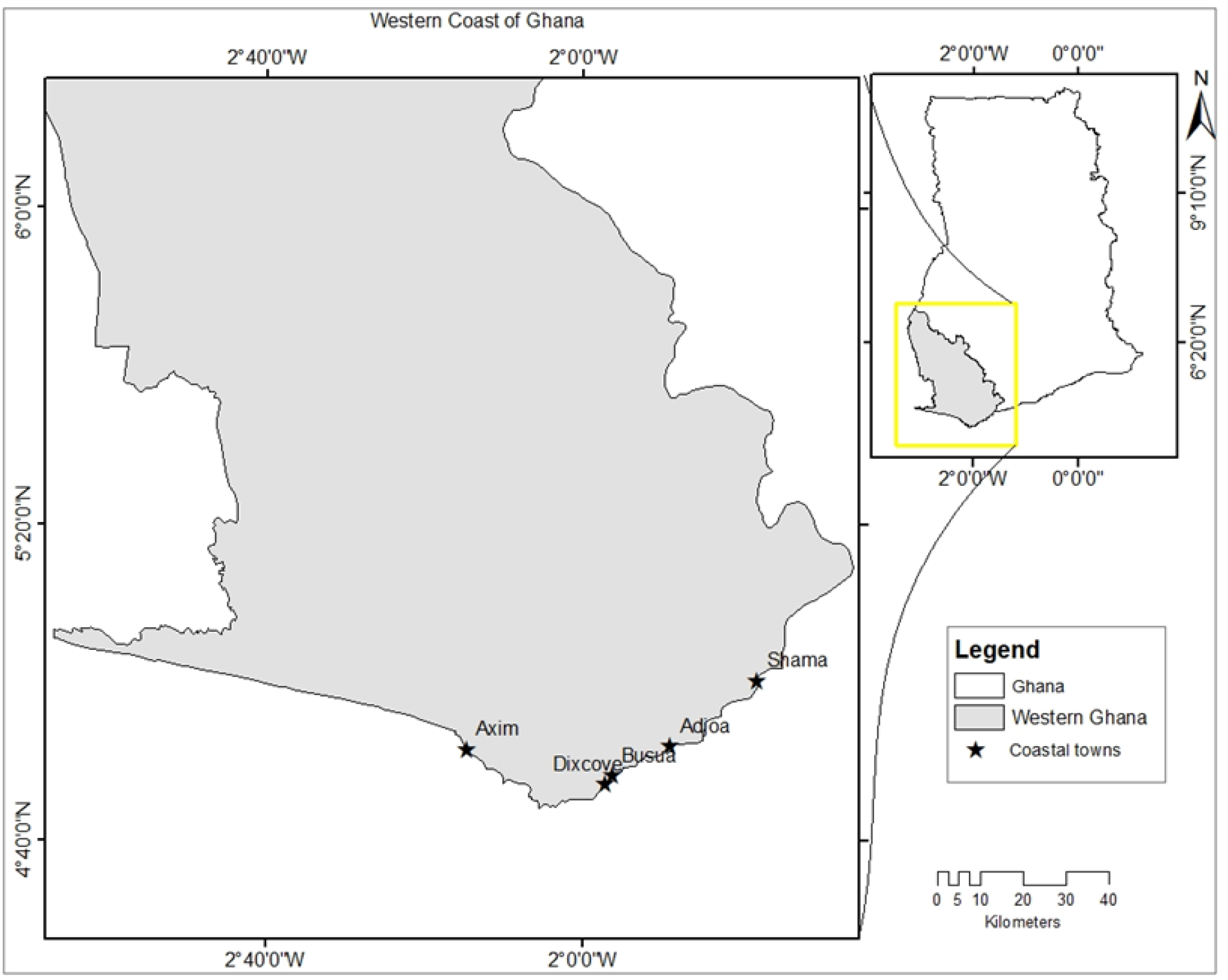
Map of Western Ghana showing the five study communities.

**Table 1:**
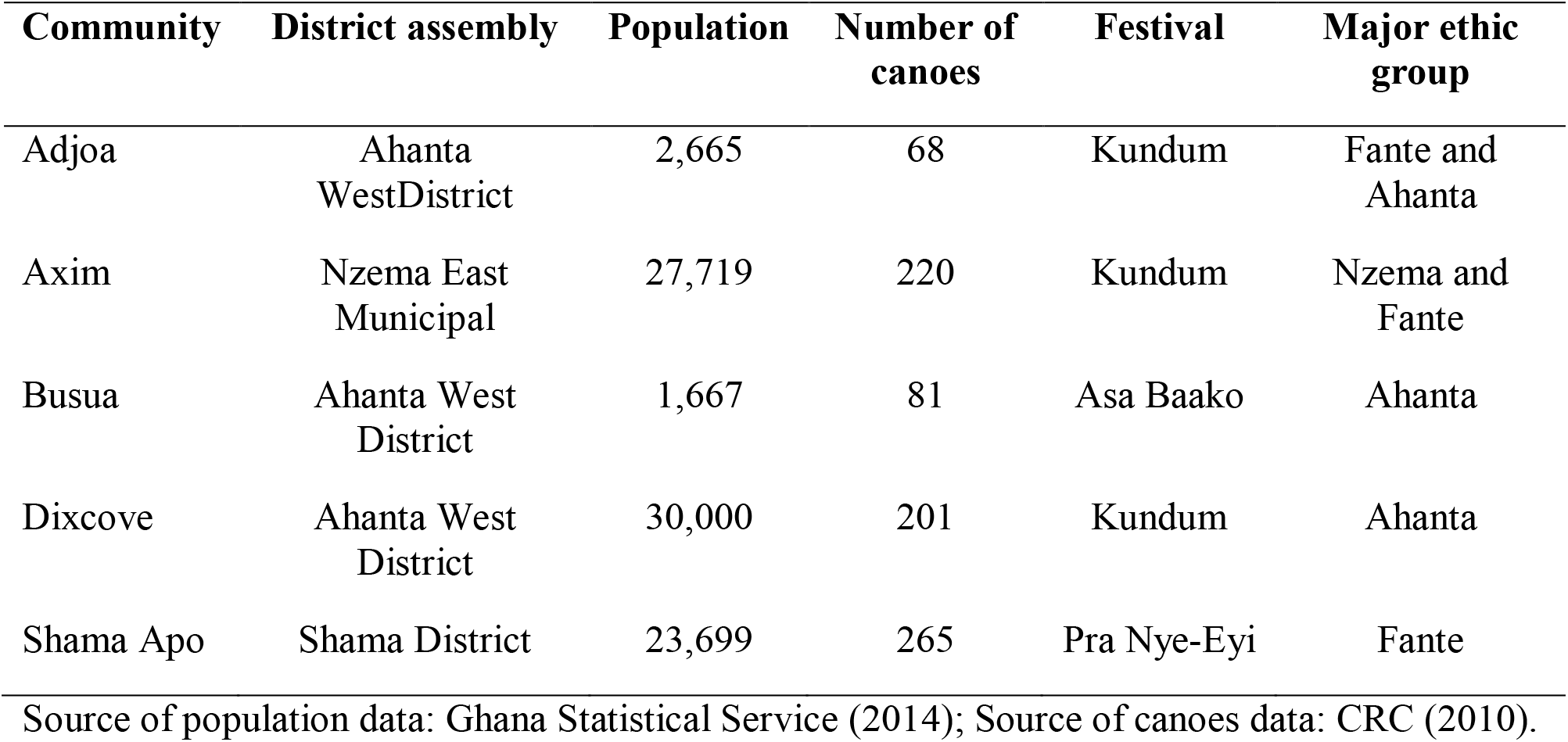
Characteristics of each study community surveyed in West Ghana.

| **Community** | **District assembly** | **Population** | **Number of**<br>**canoes** | **Festival** | **Major ethnic**<br>**group** |
| --- | --- | --- | --- | --- | --- |
| Adjoa | Ahanta<br>West District | 2,665 | 68 | Kundum | Fante and<br>Ahanta |
| Axim | Nzema East<br>Municipal | 27,719 | 220 | Kundum | Nzema and<br>Fante |
| Busua | Ahanta West<br>District | 1,667 | 81 | Asa Baako | Ahanta |
| Dixcove | Ahanta West<br>District | 30,000 | 201 | Kundum | Ahanta |
| Shama Apo | Shama District | 23,699 | 265 | Pra Nye-Eyi | Fante |

Source of population data: Ghana Statistical Service (2014); Source of canoes data: CRC (2010).

### 2.2 Data collection

Data were collected through participant observations, face-to-face interviews with 33 respondents, five focus group discussions and participatory rural appraisal methods. Data collection commenced in June 2019 and ended in July 2020. We used participant observation to gain an understanding of the main social dynamics in the study communities and became familiar with the various activities and fishing practices in the communities. The first author also participated in fishing activities in the communities, which included hauling marine megafauna ashore, butchering catch, and mending nets. He participated in all these activities with the intention of building trust with fishers in the various communities. Several meetings were held in small groups of fishers and chief fishers to explain the aim and rational behind the research and to garner their support for the research in their landing sites. The questions and interview guides were pre-tested in March 2020 in the communities of Axim and Shama, with eight respondents (four from each community).

We selected only retired and active fishers for this study because they have more direct interactions with elasmobranchs and other marine resources and are primary stakeholders in fisheries in Ghana. Further, the retired fishers have gained much experience in fishery activities and possess a wealth of retrospective knowledge on shark fisheries and operations. In contrast, active fishers possess current knowledge on fisheries, elasmobranch catch dynamics and composition. A non-probability convenience sampling approach (Alexander *et al.,* 2017) was used to select active fishers for the interview. The convenience sampling scheme also referred as availability sampling is based on the availability and willingness of respondents to participate in the interview (Newing, 2010; Naderifar *et al.,* 2017). The snowball sampling scheme was employed to track down retired fishers who doubled as canoe owners. The snow ball sampling technique was selected as referral or chain sampling is used where potential participants are difficult to find (Newing, 2010). This sampling scheme is where research participants recruit other respondents for the study (Naderifar *et al.,* 2017). These sampling schemes were employed to select respondents owing to the fact that most fishers are aware of the global controversies surrounding sharks and the fin trade, which makes it difficult to get fishers who are willing to participate in such interviews and open up to researchers.

The main data source for this research was from face-to-face interviews with respondents. A total of 33 respondents comprising 17 retired fishers and 16 active fishers were interviewed. We also interviewed three fishery officials from the Fisheries Commission to validate some key information from the fishers where necessary. Interviews were conducted at landing sites for active fishers, and mostly at the homes of retired fishers, and lasted between 60 and 80 minutes. The interviews were conducted mostly in the local languages used by each respondent and were conducted by the first author with the assistance of local volunteers fluent in English and Fante, Ahanta, Nzima or Twi. The volunteers assisted the first author in translating information from Nzima and Ahanta languages to Twi or English when necessary. The purpose of the research was explained to all respondents and their consent sought, prior to the interview. Respondents were also assured of confidentiality, and their right to omit uncomfortable questions or withdraw from the interview at any stage.

Almost all questions used for the interviews were semi-structured, which allowed the researcher to further probe interesting issues raised by the respondents. The researcher used the same questions for the interview in all five communities with some questions adapted to suit the local context. For example, some questions were specifically asked in communities where bottom-set gillnet or drift gillnet gears were used by fishers. Interviews were guided by sets of questions categorized into themes to understand the ecological factors influencing fishing operations, and changes in shark and ray stock status. Specifically, questions were designed to generate comprehensive information regarding how well fishers understand ecological factors and their influence on fishing operations. Interview questions provided an understanding of: i) bait types and how they influence the catch of sharks; ii) seasons and weather conditions; iii) the effect of lunar phase on fishing activity; iv) presence of sea birds and fish movement; v) seawater color and vi) sea currents on fishing operations. In addition, interview questions were designed to collect data on species targeted, seasons with most elasmobranch catch and changes in catch location. Changes in stock status with a focus on body sizes, species composition, and decadal changes in abundance of well-known commercially important sharks and rays since 1980, and factors causing the changes were also collected using trend analysis. This included a qualitative ranking of abundance of 10 commercial shark and seven ray species, which are well-known to fishers. Fishers were asked to characterize the abundance of each shark and ray species into one of four qualitative categories (abundant, common, depleted, and severely depleted) for each period – that is, 1980s, 1990s, 2000s, 2010s, and 2020s. Fishers were asked to quantify the abundance categories of each species using 10 stones. Seven to 10 stones indicated abundant, 5-6 indicated common, 3-5 indicated depleted, and 1-2 indicated severely depleted.

In addition, a total of five focus group discussions (one per community) were organized for both active and retired fishers at the various communities to validate the interview data. Each focus group discussion comprised six to nine participants. We slightly adapted the interview questions and used similar protocols for the focus group discussions to collect data on changes in shark and ray catch abundance, size, and composition.

### 2.3 Data analysis

Content or thematic analysis was the principal protocol used for analyzing the data. Interview notes, field records, and focus group discussion (FGD) summaries were first written in descriptive form and transcribed from spoken format to formal English. Categories from the interview data and FGD summaries were identified and aggregated manually. The data were coded and analyzed using descriptive statistics and presented in tables or figures where necessary.

For the decadal changes in abundance of commercially important shark and ray species, we assigned the ordinal abundant categories (i. e. abundant, common, depleted, and severely depleted) a category weight of 1, 0.75, 0.5, and 0.25 respectively. The weighted score for each species in a given decade was then calculated using the formula;

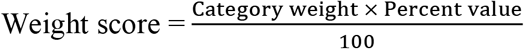

Where percent value is percentage of all respondents reporting abundance, common, depleted, and severely depleted for a species in a given decade.

## 3. Results

### 3.1 Demographic characteristics of respondents

All 33 respondents were males, as women are barred from fishing due to traditional beliefs. Of the 33 respondents, 52% (n = 17) were retired fishers who doubled as canoe owners and 48% (n = 16) were active fishers. The mean age of respondents was 45.9 ± 15.0. Fishing experience of both retired and active fishers ranged from 6–58 years. Many of the respondents (64%) had no education. Most of them belong to the Fante ethnic group (Table 2).

**Table 2:** Demographic characteristics of respondents from each community surveyed in West Ghana

| Variables | Adjoa | Axim | Busua | Dixcove | Shama | Sum/<br>average |
| --- | --- | --- | --- | --- | --- | --- |
| No. of retired fishers | 2 | 4 | 2 | 4 | 5 | 17 (52%) |
| No. of active fishers | 3 | 4 | 3 | 1 | 5 | 16 (48%) |
| Total no. of respondents | 5 (15%) | 8 (24%) | 5 (15%) | 5 (15%) | 10 (30 %) |  |
| **Mean age** | 42.0 ± 16.8 | 43.0 ± 12.2 | 42.0 ± 18.3 | 54.6 ± 11.6 | 47.7 ± 16.8 | 45.9 ± 15.0 |
| Minimum age | 26 | 22 | 20 | 42 | 27 |  |
| Maximum age | 64 | 56 | 62 | 65 | 70 |  |
| **Educational level** |  |  |  |  |  |  |
| Illiterate | 4 | 5 | 4 | 3 | 5 | 21 (64%) |
| Junior school | 0 | 1 | 1 | 0 | 2 | 4 (12%) |
| Senior High | 1 | 0 | 0 | 2 | 1 | 4 (12%) |
| Tertiary | 0 | 2 | 0 | 0 | 2 | 4 (12%) |
| **Years of fishing experience** | 23.8 ± 12.0<br>(11 - 41) | 28.3 ± 12.4<br>(10 - 43) | 27.2 ± 17.6<br>(16 - 50) | 33.6 ± 14.2<br>(20 - 50) | 31.4 ± 18.0<br>(10 - 58) | 29.2 ± 14.7<br>(6 - 58) |
| **Ethnic groups** |  |  |  |  |  |  |
| Fante | 0 | 6 | 0 | 3 | 10 | 19 (58%) |
| Ahanta | 4 | 1 | 4 | 0 | 0 | 9 (27%) |
| Nzima | 1 | 1 | 1 | 2 | 0 | 5 (15%) |

% represents percent of total respondents for each variable

### 3.2 Fishing gears used in the communities

The study sites have a multi-gear fishery. However, sharks and rays are caught with two major gear types; drift gillnets complemented with longlines as well as bottom-set gillnets. Baited hooks are deployed as secondary longline gears, which are set alongside the drift gillnet in the same fishing grounds and directions. The gears used by fishers were dependent on the size of canoe, fishing grounds (oceanic or coastal) and in many instances the finances of the canoe owners. All fishers interviewed from Shama, Dixcove, and Axim used drift gillnets complemented with longlines gear (n=23), while fishers from Adjoa and Busua used bottom-set gillnets (n=10) for their fishing operations. Both gear types were made of monofilament fishing line. The gears are deployed and retrieved manually by the fishers.

### 3.3 Learning, understanding and use of ecological variables in fishing operations

Fisher‟s knowledge regarding fishery activities and their practices generally stem from older generations and their continued interactions with their surroundings, especially during fishing trips. Half the fishers (48%) learned fishing practices and ecological factors from their families (grandfathers, fathers, and uncles), as apprentices in a crew (16%), or in some cases through observation (36%) in their communities. These ecological clues have been applied over the years to enhance fishing operations as well as maximize fisher catch. Fishers were able to identify and name species they target, their composition, and abundance. They rely on local ecological knowledge to assess potential fishing locations, timing, and protocols for their fishing activities. The interviewed fishers described six categories of ecological factors they use as clues and their understanding of how these clues affect their mode of operations on the sea. The stated clues are subdivided into two main sections, which are presented below.

#### 3.3.1 Fishers knowledge regarding timing and protocols of fishing

Knowledge about periods and practice of fishing gave fishers opportunities to know when and how to fish. Such knowledge includes seasons and weather conditions, lunar phase, and bait type.

##### 3.3.1.1 Seasons and weather conditions

All interviewed fishers agreed that knowledge about bumper and lean seasons as well as the reproductive seasons for pelagic and demersal species are indispensable in preparing and targeting particular fish taxa. According to them, harvesting seasons for both pelagic and demersal species vary considerably. Most drift gillnet operated fishers (87%) stated that the harvesting season for sharks, billfishes, and most types of tunas is from July to September. The lean seasons for these species are December to February. Fishers recounted that the breeding season for most tuna including Frigate Mackerel (*Auxis thazard*), Skipjack (*Katsuwonus pelamis*), Yellowfin Tuna (*Thunnus albacares*) and Bigeye Tuna (*Thunnus obesus*) is from May to July and that of sharks is March to June. This is the period they mostly catch and see neonates. Most of the interviewed bottom-set gillnet fishers recounted that the harvesting season for rays, especially Stingrays (*Fontitrygon* spp), Cassava Fish (*Micropogonias undulates*), and Threadfin Fish (Polynemidae) is from August to October.

Considering weather conditions, 82% of fishers reported that the major rainy season (May to July) was not a good period for fishing. This period, they recounted, is characterized by high fog, which makes navigation difficult, and the heavy rainfall is normally accompanied with strong storms, which can capsize canoes and displace fishing nets. Many of the respondents (73%) mentioned the minor rainy season (August to November) and dry season (November to March) as the best periods for fishing.

##### 3.3.1.2 Lunar phase

The majority of fishers (94%) reported that the moon affects their ability to catch fish species of all kinds. These include both large pelagic species like Shark, Swordfish, Sailfish, and Tuna, and demersal species like Stingray, Guitarfish, and Anchovy in large quantities. Fishers reported that a full moon makes the water more transparent for these pelagic species to see and avoid the gears. To fishers, the best periods to go fishing are when there is partial or no moon. This, they said, gives them the opportunity to get substantial catch because of the opaqueness of the water.

##### 3.3.1.3 Bait type

All the interviewed fishers‟ operating with drift gillnets, which are complemented with longline gears, recounted that the type of bait used has a significant influence on the catch of large pelagic fish species. They mostly use freshly caught Dolphin, Flying Fish, Tuna, and Sardinella as bait (Table 3). Dolphins were regarded as the best and most effective bait for catching sharks by all respondents, followed by Flying Fish. To fishers, the bright and shiny skin of dolphins attracts sharks more quickly than any other bait. Further, all respondents reported that both Dolphin and Flying Fish have a powerful odor and are characterized with high oil and blood content, which are good attractants for sharks. Dolphins are mostly bought from Dixcove in the Western Region, Tema in the Greater Accra Region, and Apam in the Central Region. Flying Fish and Tuna are mostly harvested by the local fishers operating with drift gillnets, while Sardinella are caught by the fishers using ring net gears. The respondents mentioned that in desperate situations where all these baits are unavailable, they mostly rely on cold-store Sardinellas, Herrings, Beef, and Pork as baits.

**Table 3:** Species used as bait in drift gillnet fisheries in the study communities of West Ghana

| Species | Common name | Scientific name | Local name | No. of respondents |
| --- | --- | --- | --- | --- |
| Dolphins | Atlantic Bumpback | *Sousa teuszii* | Etwui | 23 (100%) |
|  | Common Bottlenose | *Tursiops truncates* |  |  |
|  | Oceanic Dolphins | *Stenella* spp |  |  |
| Flying fish | Unspecified | Exocoetidae (family) | Epoanomaa | 19 (83%) |
| Tuna | Frigate Mackerel | *Auxis brachydorax* | Apaku | 14 (61%) |
|  | Atlantic Little Tuna | *Ethynnus alleteratus* | Apakuwona |  |
|  | Chub Mackerel | *Scomber japonicas* | Akowona |  |
| Sardinella | Round Sardinella | *Sardinella aurita* | Eban | 8 (35%) |
|  | Flat Sardinella | *Sardinella maderensis* |  |  |

#### 3.3.2 Fisher knowledge regarding oceanographic conditions

Knowledge of oceanographic conditions provided fishers with clues of fishing locations. These factors included sea birds and fish movement, color of seawater, and sea current.

##### 3.3.2.1 Seabirds and fish movement

Most interviewed fishers (88%) reported that seabirds are good indicators for the presence of fish and potential fishing areas. Fishers reported that some birds have the ability to see deep below the water surface to hunt and feed on smaller fishes such as Anchovies and Sardinellas, which invariably serve as foods for large pelagic species including sharks and billfishes. Hence, the presence of a large flock of birds hovering close to the surface of the seawater and dipping their beaks in the water is an indication that there are fish, and the fishers set their nets along that particular area. Fishers further stated that knowledge regarding the movements of seabirds combined with an understanding of the rippling effect of seawater and fish movement are important clues used for identifying a prospective fishing location. According to respondents, a faster movement of fishes from a particular area, or when they are seen jumping out the water, gives an indication that setting a net in that location may not yield a good catch. However, fishes moving in circular motions give an indication to fishers that they are stable within that particular location and fishers therefore have high possibilities of getting abundant catch should they set their nets there.

##### 3.3.2.2 Seawater color

Interviewed fishers reported five different colors of seawater – deep blue, light blue, yellow, red, and light green (Table 4). Deep blue color, referred to as “Adomnsuo” in the local dialect, literally means “divine water”, and was listed as the preferred color by all respondents (100%). It was followed by light blue/sea blue (91%). Yellow color (33%) was the least preferred seawater color for fishers and occurs intermittently at any period within the year. Fishers stated that Cassava Fish, Herring, and Bill Fish are mostly caught in yellow water whenever it occurs. Most respondents (78%) stated that deep blue water occurs unpredictably once a year between July and October at any location in the sea, and does not last for more than two weeks. Deep blue water provides bountiful catch of all kinds of fish to both drift gillnet and bottom-set gillnet fishers. Light green water provides good conditions for larger fish such as sharks, tunas, and bill fishes, and according to fishers can occur randomly at any period within the year. Red color occurs intermittently in June to July and is mostly associated with tunas (Frigate Mackerel, Skipjack), Guitarfishes, Whiprays, Cassava Fish, and Octopus. Light blue water occurs all year round whenever they go fishing and can get almost all kinds of pelagic and demersal fishes.

**Table 4:** Fishers‟ perception of colors of seawater and associated favored fish species

| **Water color** | **Local names** | **Most favored species** | **No. of respondents** |
| --- | --- | --- | --- |
| Deep blue | Adomnsuo | All kinds of fish, both demersal and pelagic | 33 (100%) |
| Light blue | Sea blue | All kinds of fish, both pelagic and demersal | 30 (91%) |
| Light green | Green water | Sharks, Tuna, and Bill Fish | 24 (73%) |
| Red | Red water | Frigate Mackerel, Skipjack, Guitarfishes, Whip Rays, Cassava Fish, and Octopus | 19 (58%) |
| Yellow | Yellow water | Cassava Fish, Herrings, and Bill Fish | 11 (33%) |

##### 3.3.2.3 Sea current

Most respondents (85%) reported that sea currents have a significant impact on fishing. Strong currents are perceived to obstruct the movement of fish and further carry them away from a particular area, while slow currents provide good conditions for fish to stay in one area. Fishers described the two main current directions they rely on while fishing on the sea – eastward and westward currents. Fishers stated that the eastward currents are mostly slow and provide favorable conditions for fish to stay at a particular area and is thus best for catching more fish. The westward currents were regarded as very strong and tend to transport fish away from the area.

### 3.4 Catch composition and abundance of target species

Fishers reported that they target multiple species in response to the seasonal changes and fluctuating abundance of most fish species. All interviewed fishers stated that they fish all year round and the number of trips per annum is mostly dependent on the financial capacity of the canoe owners and availability of premix fuel. Most respondents (91%) stated that July to September was their ‘bumper’ season when the greatest catches were taken. Fishers were quick to add that their fish stocks have severely depleted presently and that there is no-longer a guarantee of getting bumper harvests in these periods. All fishers are now generalists who target several species of fish. Drift gillnet fishers target large pelagic species but occasionally catch smaller ones, while the bottom-set gillnet fishers target demersal fishes and occasionally catch some pelagic species (Table 5). When asked about the most uncommon species of fish, most drift gillnet fishers stated that all shark species, with the exception of Blue Shark (*Prionace glauca*) (87%), Devil rays (*Mobula* spp) (78%), and Marlins (Istiophoridae) (74%), are now becoming increasingly difficult to catch. Many interviewed fishers (65%) further listed tunas (Frigate Mackerel, Skipjack, Yellowfin, and Bigeye Tuna) as species dominating their catch. However, most bottom-set gillnet fishers (80% of 10 respondents) stated guitarfishes and herrings as the rarest fish they catch. Fishers stated Cassava Fish and rays as the most common species they catch.

**Table 5:**
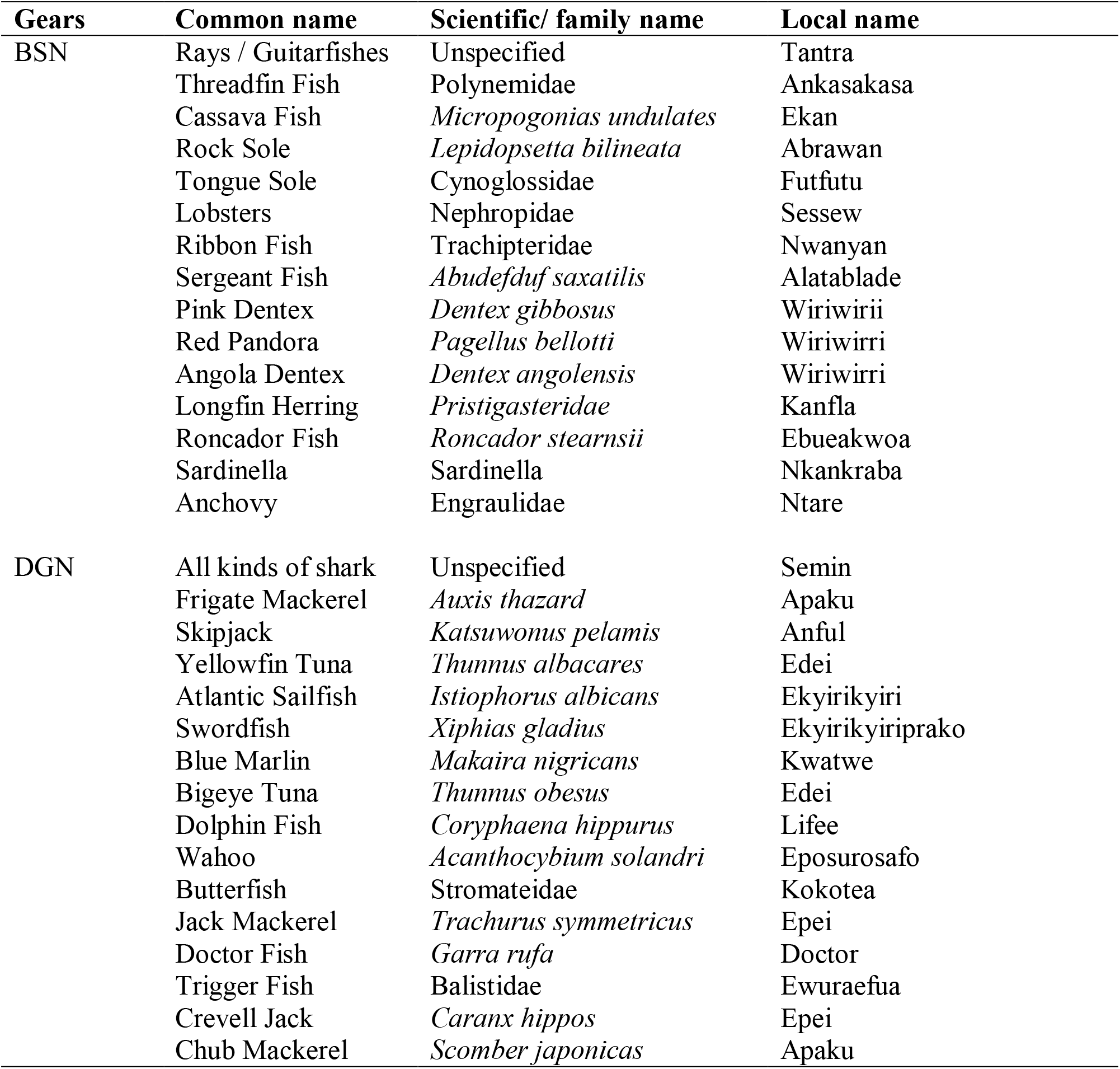
Fish species targeted in bottom-set gillnet (BSN) and drift gillnet (DGN) gears in the study communities.

### 3.5 Changes in elasmobranch-specific catch composition and sizes

Fishers in the study communities have a long standing history of catching sharks since the early 1970s. Retired fishers recounted that they used to target small pelagic and other demersal species with bottom-set gillnets and paddle canoes; setting their nets in coastal zones and retrieving them the next day. Fishers recounted that there was incessant destruction of their gears by sharks and other bigger fishes and this motivated some of them to develop nets to suit the sizes of these big fishes in the early 1970s. Fishers further reported that sharks and rays became a major target in the 1980s, when Ghanaians residing in the neighboring West African countries including Guinea, Gambia, Senegal, and Mali returned home and started trading in shark fins, which afforded them much profit. A retired fisher in Shama recounted that:

> “When we started fishing about 50 years ago, our intention was never to catch sharks. Sharks were not valuable at all. Many fishers were not actually involved in shark fisheries. Most fishers were interested in Tunas, Anchovies, and Sardinellas and were catching more in the coastal habitats. I started targeting sharks when I got much profit on my first fin trade in the late 1980s.” (Retired fisher, Shama, 06/2020).

Interviewed fishers recalled their knowledge about elasmobranch species they catch, their abundance, and composition. The most well-known and valuable species to respondents were Blue Shark (*Prionace glauca*) and Devil rays (*Mobula* spp), which were mentioned by all drift gillnet fishers, while Whale Shark (*Rhincodon typus*) was mentioned the least by respondents (17%) (Table 6). Fishers reported that Blue Shark were the most common shark species caught in drift gillnet gears. This assertion was confirmed by the participants in the focus group discussions as well as during participant observation.

**Table 6:**
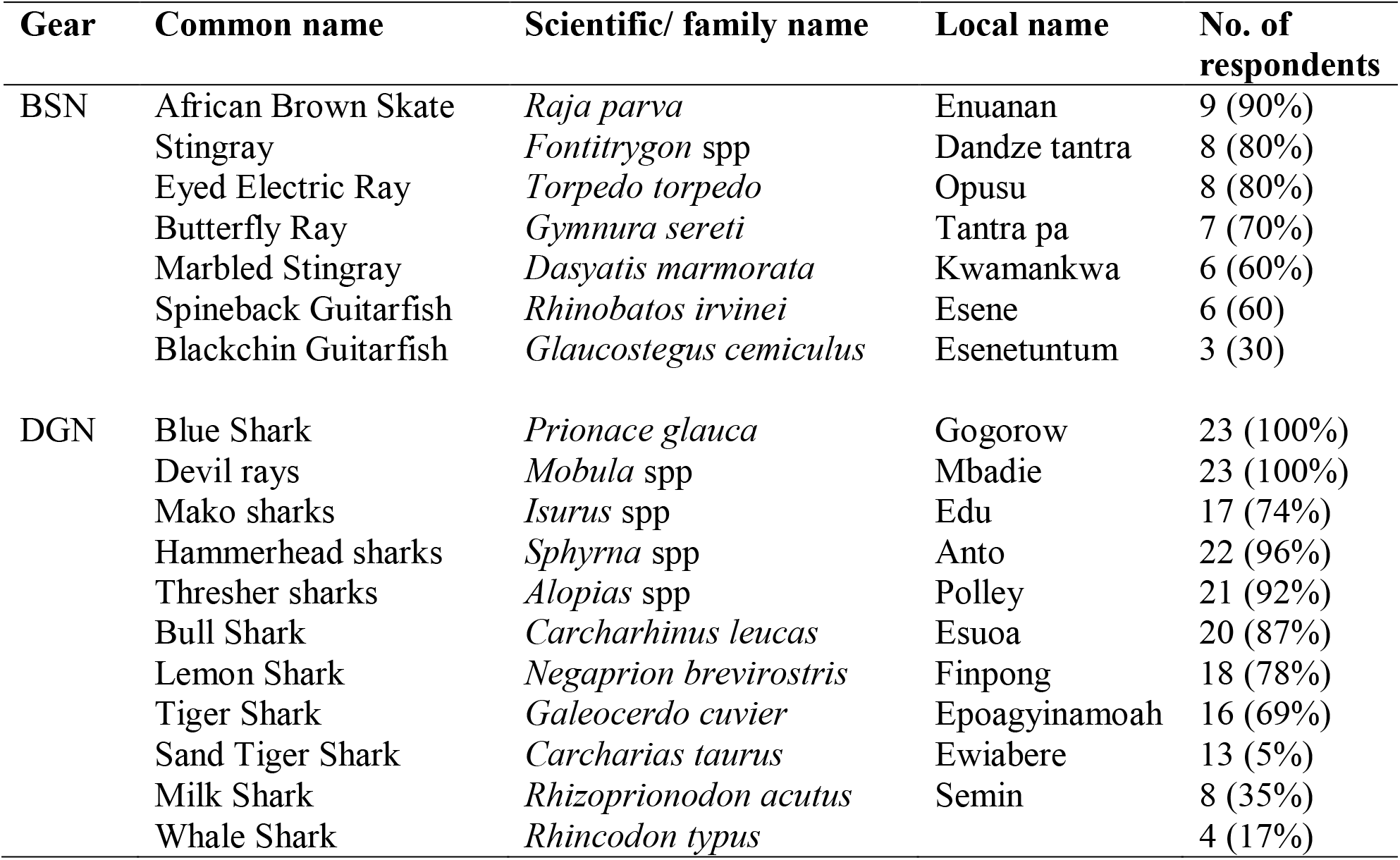
Shark and ray species listed by drift gillnet (DGN) and bottom-set gillnet (BSN) fishers in the study communities.

With reference to bottom-set gillnet fisheries; fishers were able to reveal knowledge about seven rays they caught in their gears. These included African Brown Skate (*Raja parva*, 90%), Electric Ray (*Torpedo torpedo,* 80%), and Stingrays (*Fontitrygon* spp, 80%), while Blackchin Guitarfish (*Glaucostegus cemiculus*, 30%) was the least caught ray species (Table 6). Most respondents (70%) stated African Brown Skate and Stingrays as the most common ray species caught. Focus group discussions and participants’ observation affirmed that these species were indeed relatively common catch in bottom-set gillnet gears.

When asked about the best seasons to catch sharks, fishers gave a wide range of answers. This is partially due to the fact that fishing is done all year round. Only 39% of fishers reported that the best seasons for catching sharks is from October to December, especially immediately after the bumper season (July to September). One fisher stated that he catches more sharks in November because they travel farther to the oceanic zone during this period as most sharks are found in deep sea. In response to the question “which location is best to catch more sharks or rays”, over 96% of fishers operating with drift gillnets stated that oceanic habitats are where they catch more sharks. Bottom-set gillnetters fish only in coastal habitats and reported that they catch all their ray species in this particular habitat. Most fishers (91%) reported that the areas they used to fish have changed significantly and that they have to travel farther to reach their fishing grounds. The respondents attributed light fishing (88%) and foreign vessels (91%) as the reasons for depleting fish stocks.

In addition to assessing the species composition of elasmobranch catch, we asked fishers to state the species of shark and ray that are now very uncommon in their catch. Lemon Shark (*Negaprion brevirostris*, 83%), Tiger Shark (*Galeocerdo cuvier*, 78%), Thresher sharks (*Alopias* spp, 73%), Hammerhead sharks (*Sphyrna* spp, 65%), Bull Shark (*Carcharhinus leucas*, 61 %), and Sand Tiger Shark (*Carcharias taurus*, 54%) were most frequently stated by fishers (n=23) as uncommon species. Most fishers have never caught or observed Whale Shark and only two had done so in more than a decade. Participants in the focus group discussions confirmed that Lemon Shark, Tiger Shark, Hammerhead sharks, and Bull Shark were the rarest species caught presently. An active fisher stated that:

> “Lemon Shark and Hammerhead sharks have always been very rare even though their fins are valuable. Our hopes have always been dashed anytime we aspired to catch some. It is 11 years now since I caught a Lemon Shark.” (Active fisher, Dixcove, 07/2020)

Most of the interviewed bottom-set gillnet fishers recalled that guitarfishes were the rarest species of fish they catch. All respondents had never seen or caught sawfishes (*Pristis* spp), with the exception of a retired fisher who claimed he saw one specimen 16 years ago at Apam landing site in 2004. Most fishers (76% of 33) stated that the sizes of sharks and rays they used to catch have decreased over the years. Respondents recollected reductions in the sizes of sharks since the 2000s and then rays since the 2010s. Similarly, when asked about the observed changes in the abundance of sharks and rays, most fishers (85%) stated that the number of sharks and rays they caught have significantly declined now compared to when they began fishing. Only two fishers reported that they catch more sharks now than when they started fishing, but this is because they have increased their number of nets and hooks deployed in the waters. Most fishers also expressed concern about the reduction in composition of sharks they catch. Fishers stated that they catch less shark or ray species (73%) and less valuable sharks or rays (67%) now as compared to previous years when they started fishing. Focus group discussions confirmed that shark and ray species composition have changed over the years. The focus group discussions also confirmed that Stingrays, Devil rays, Blue Shark, and African Brown Skate have the least valuable meat although these species dominate their catch. An active fisher narrated:

> “We used to get guitarfishes and I mean the bigger ones in large quantity. These days, even getting one individual in a fishing trip is a problem. The ones we get are not big enough and we do not even get fins from them to sell.” (Active fisher, Adjoa, 06/2020)

Fishers who had noticed changes in their catch were asked the reasons for this occurrence. The major reason cited by fishers (66% of 32 fishers) for the changes in sizes, abundance, and composition of sharks was the practice of Illegal, Unreported and Unregulated fishing (IUU) (Figure 2). Fishers were also increasingly concerned about the activities of foreign vessels on the sea, which are invading their territories and engaging in illegal fishing practices as well as catching more by-catch that exceeds their quotas sanctioned by the laws of Ghana. Competition with foreign vessels was mentioned by 63% of fishers as being a possible cause of decline in abundance and composition of sharks and rays. Fishers attributed the reduction in shark species abundance and sizes to the escalating rate of light fishing and dynamiting, which are depleting the smaller fishes such as Anchovies and Sardinellas. Fishers stated that these species are food for sharks and once they are depleted sharks will also travel far into the deep and oceanic waters to find other sources of food. Most fishers believed that the smaller sharks that cannot travel far remain and feed on the remaining food and that is the main reasons why they get smaller sized sharks and rays these days. Another reason was overfishing, which was attributed to increasing number of canoes and nets in the sea (64%).

**Figure 2:**
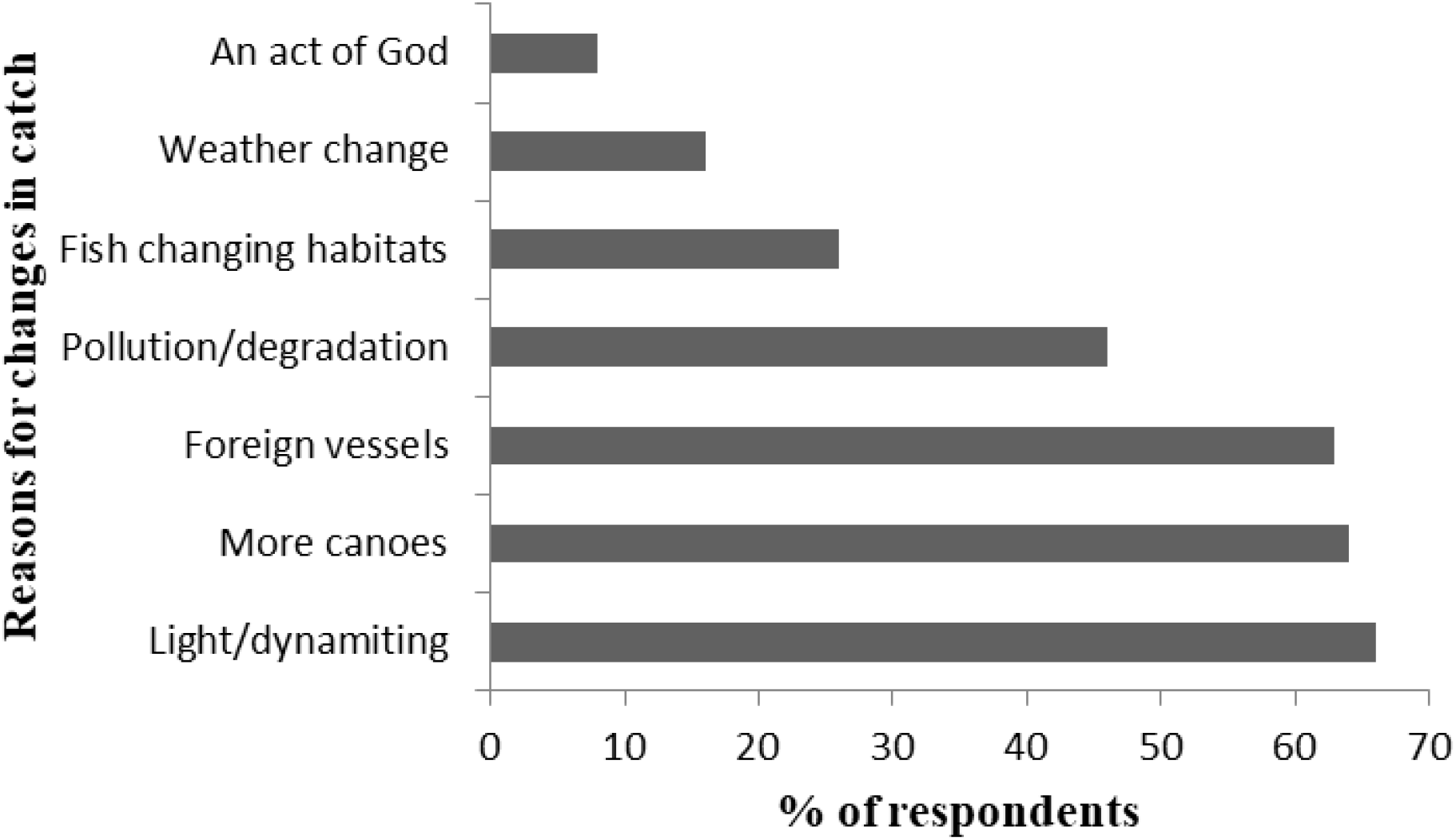
Perception of fishers on reasons for changes in shark and ray species catch composition. The figure contains multiple responses

### 3.6 Changes in abundance of elasmobranch since 1980

We used trend analysis to rank the abundance of each genus or species of shark and ray from 1980 to 2020 (Figures 3 and 4). In general, most shark and ray species were abundant in 1980 but have been severely depleted as of 2020. Most shark and ray species got depleted in the 2010s with a weighted score less than 0.75, while other species started depleting in the 2000s. Shark and ray species that were depleted in the 2010s were *Sphyrna* spp, *Carcharhinus leucas*, *Carcharias taurus*, *Dasyatis marmorata*, *Fontitrygon* spp, and *Rhinobatos irvinei*. Species such as *Alopias* spp, *Galeocerdo cuvier, Glaucostegus cemiculus, Isurus* spp, and *Negaprion brevirostris* were depleted as early as in the 2000s. All shark species (except *Negaprion brevirostris*, *Alopias* spp and *Galeocerdo cuvier*) and all ray species were abundant in 1980s. While no species of shark and ray was abundant from 2010s, *Prionace glauca* and *Mobula* spp were reported to be common by the interviewed fishers.

**Figure 3:**
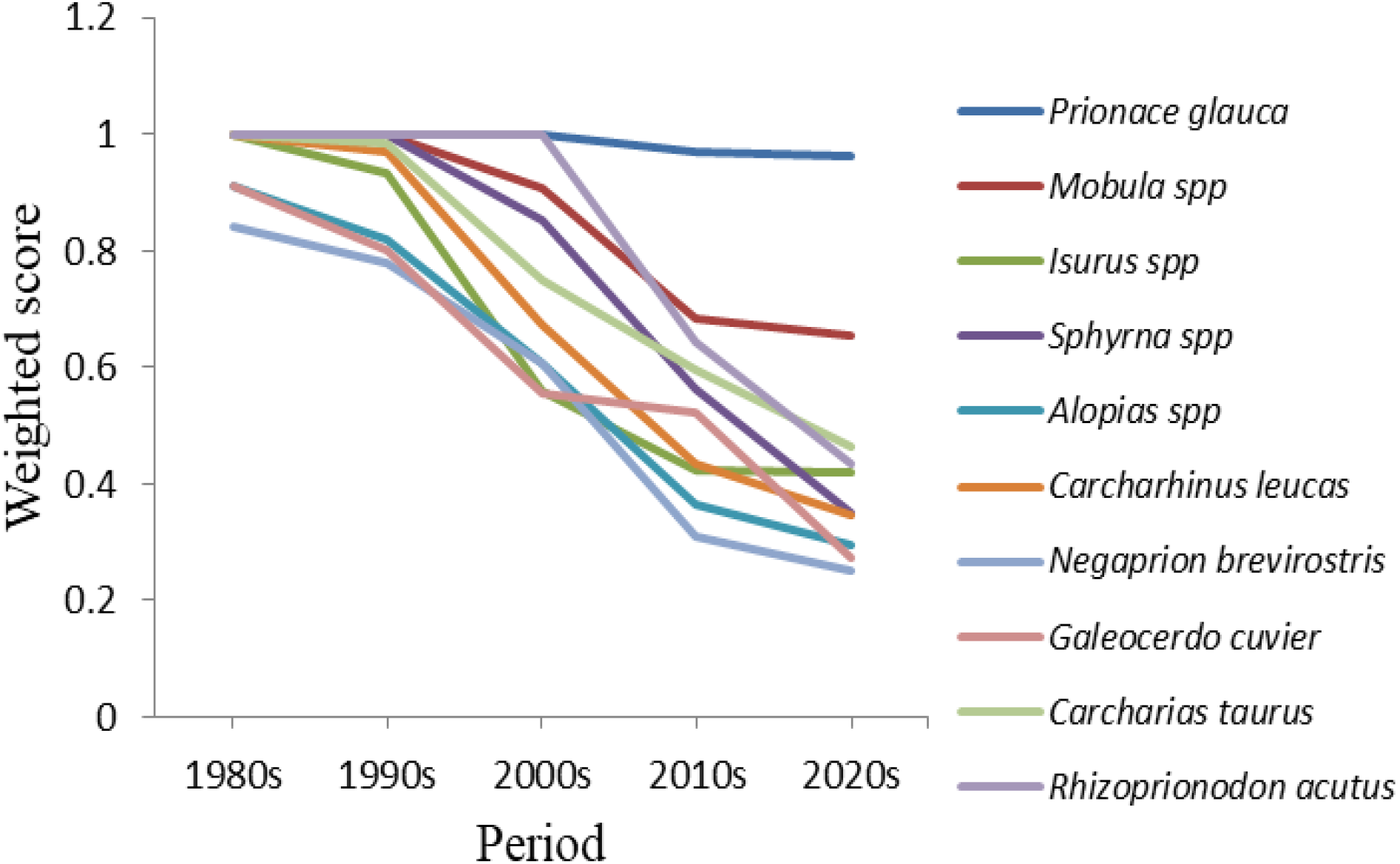
Perceived changes in abundance or otherwise of sharks by drift gillnet fishers (n=23) from 1980 to 2020

**Figure 4:**
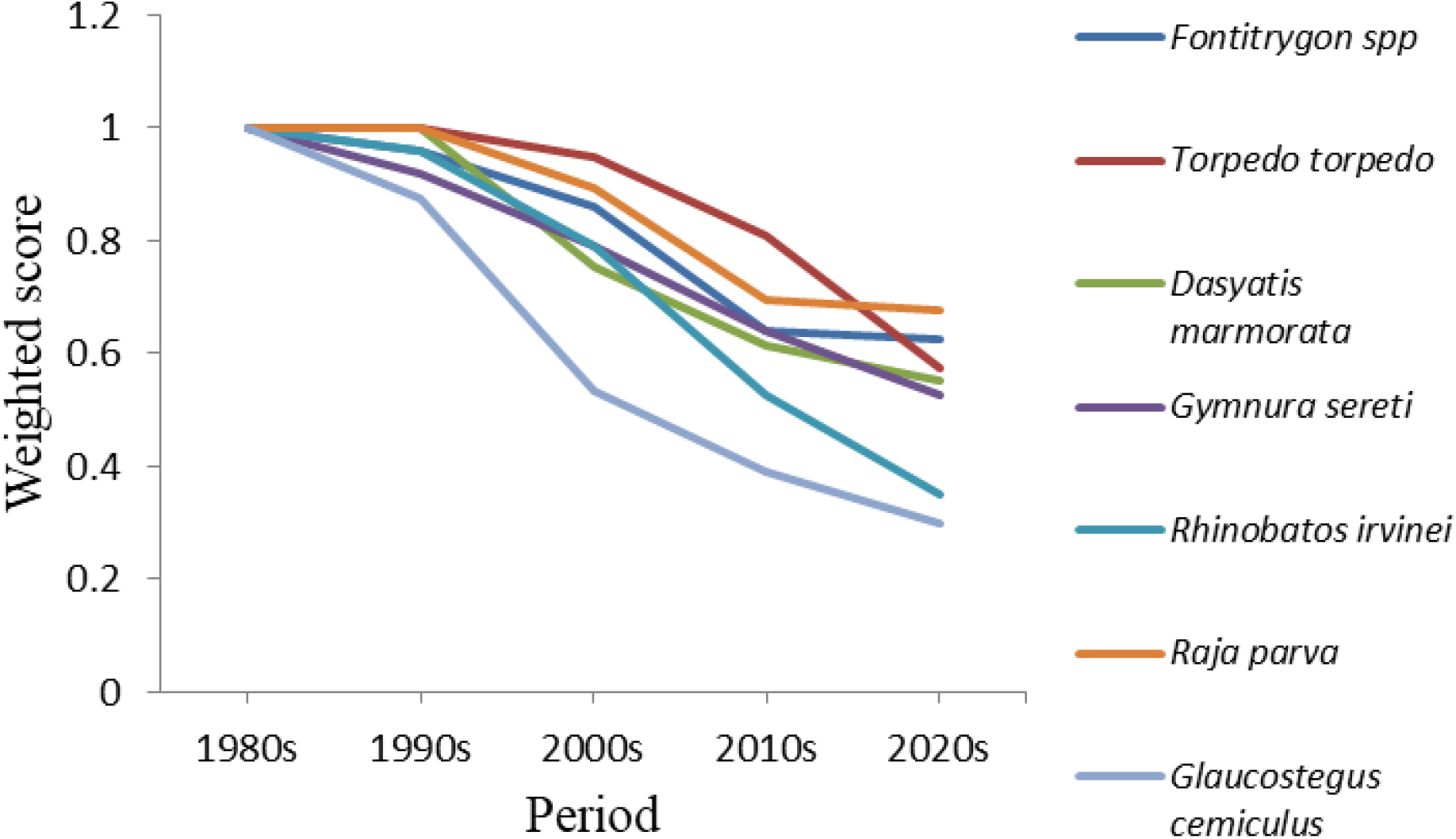
Perceived changes in abundance or otherwise of rays by bottom-set gillnet fishers (n=10) from 1980 to 2020

## 4. Discussion

Overall, we found that as of 2020s shark and ray species catch has been severely depleted in Western Ghana, with the exception of *Prionace glauca* and *Mobula* spp, which were reported to be common by fishers. Half the fishers learned fishing practices and ecological variables from their families and these six ecological features have been successfully applied over the years to maximize fisher catch. Our findings that fishers are knowledgeable about the diverse distribution and abundance of sea life from season to season based on habitat, weather, lunar phase, currents, and other environmental factors agree with the findings of Johannes and Neis (2007). These environmental variables that repeatedly occur are invariably mastered by fishers based on learning and experience, which provide guidance to facilitate and improve the effectiveness of their fishing operations while at sea (Silva, 2005). Some local ecological variables described by fishers in the study communities are consistent with published scientific data. These include the use of birds as fish indicators (Gomes *et al.,* 1998; Vlietstra, 2005), harvesting and spawning seasons of large pelagic species (Oxenford, 1999) and the influence of the Amazon on seawater color in the eastern Caribbean (Gomes *et al.,* 1998). In addition, several studies have revealed how fisher’s understanding of ecological and biological variables influence their fishing operations. For example, fishers in Patos Lagoon, Brazil have observed that Silver White Croaker (*Pennahia argentata*) catches are hindered by the brightness or fullness of the moon and attributed this phenomenon to the active nature of these species during this lunar phase (Kalikoski and Vasconcellos, 2007). The identified local ecological variables of fishers in the present study are consistent with those of fishers from Gouyave, Eastern Caribbean (Grant and Berkes, 2007). The Gouyave fishers described the presence of seabirds, seawater color, weather conditions, and sea current, amongst other variable, as ecological factors which influence their longline fishing operations (Grant and Berkes, 2007).

The claim by fishers that sharks and rays became a major target in the 1980s corroborates the study of Clarke (2005) who reported that elasmobranchs were primarily caught as bycatch until the rise in international demand for their products, particularly fins, in the mid-1980s. This incentivized many coastal communities to target sharks. The rise in fin trade was largely attributed to the reforms of the economy and increasing rate of the wealthy population in China, which led to the reintroduction of shark fin soup in the 1980s (Cook, 1990; Buckley and Hile, 2007; Fabinyi, 2012). This was also the period when marine biologists and conservationists started to realize that many shark populations were under severe threat (Manire and Gruber, 1990).

The results of this present study show that fishers have noticed changes in abundance of *Negaprion brevirostris*, *Galeocerdo cuvier,* and Rhinobatidae which had the most valuable meat and fins in 2000s. A strong trend in fishers responses indicates a depletion of *Dasyatis marmorata*, *Sphyrna* spp, *Carcharhinus leucas* amongst others in the 2010s, while *Alopias* spp, *Galeocerdo cuvier, Glaucostegus cemiculus, Isurus* spp, and *Negaprion brevirostris* were reported to be severely depleted as early as the 2000s. It may be possible that most shark species reported to have changed in abundance in the 2000s were already showing signs of depletion in the 1990s, but most fishers failed to notice that. This is because fishers reported that in the mid-1990s to 1998, fishing pressure on sharks was remarkably reduced, as most fishers using large canoes with drift gillnet gears migrated to Liberia, Gambia, Benin, and Cote d’Ivoire to fish in their waters. A focus group discussion confirmed that most fishers had already started noticing depletions in a number of shark species in the early 1990s and were forced to move to countries where they could get abundant shark catches. Thus, the few fishers who remained and were fishing in Ghanaian waters in the 1990s had abundant catch. This partly explains why fishers reported most sharks to be abundant in this period. Migrant fishers returned home and started their fishing operations in Ghanaian waters between 1999 and early 2000 when they noticed that the shark species in the other countries were becoming increasingly depleted. This is the period when fishing pressure intensified on sharks and other pelagic species. Fishers catch in the present year (in 2020) mostly encompassed species such as *Prionace glauca*, *Mobula* spp, and *Raja parva* with other species remarkably declined. The results agree with those of Jaiteh *et al.* (2017), where Indonesian fishers also reported a decline of 16 shark and ray species including *Sphyrna* spp, *Galeocerdo cuvier* and Rhinobatidae between 1993 and 2013 in Eastern Indonesia. Similarly, Ainsworth *et al.* (2008) also found that fishers reported a steep decline in shark catches in Raja Ampat, Indonesia, between 1990 and 2000. In West Africa, trend data are lacking, but there is evidence of severe declines of Wedgefishes across its range. Wedgefishes was moderately abundant across its former range in 1960 in West Africa, but declined dramatically thereafter (Kyne *et al.,* 2020). Similar to our findings, in southern areas of West Africa (Cameroon to Namibia) qualitative evidence seems to be stronger for extinction of Sawfishes across their range than against, although uncertainty because of the absence of data in this subregion (Fernandez Carvalho *et al.,* 2013). Further, reviews of recent status of sawfishes from Mauritania to Republic of Guinea in West Africa revealed that sawfishes, *Pristis pectinata* and *Pristis pristis* which was referred as *Pristis microdon* were relatively common in the past but are now rarely caught or observed (Ballouard *et al.,* 2006; Robillard and Séret, 2006).

An indication of overfished stocks is the decline in abundance and sizes, as well as changes to the composition of shark and ray species fishers used to catch. Although there are no retrospective data to confirm this claim, which calls for caution in interpreting the findings, the fact that fishers are travelling longer distances to fishing grounds than before is an indication of decline in catch of sharks and rays in the study communities. Other parallel studies have demonstrated that fishers traveling far distances to offshore zones are signs of overfishing and stock decline. For example, in China, a similar scenario has been reported with fishers moving further offshore and intensifying their fishing efforts, although their catch remains virtually the same (Lam and Sadovy de Mitcheson, 2011). Similarly, Wing and Wing (2001) found that the mean trophic level of reef fishes declined from the early to late occupation on each of the five studied Caribbean islands and further suggested that populations of reef fishes adjacent to occupation sites on these islands were heavily exploited in prehistoric times.

The perceived decrease in abundance encompasses species that were mostly exploited by fishers and are regarded as commercially important in both the drift gillnet and bottom-set gillnet fisheries. Consequently, species such as Common Smooth-hound (*Mustelus mustelus*) and Spiny Butterfly Ray (*Gymnura altavela*), among others, which were occasionally observed in landing sites during participant observation, were not familiar to most fishers and for that matter were not easily identified even when photo illustrations were shown to them. This may be partially explained by the rare nature of these species in fishers catch or because these species are not particularly valued by fishers, as some of them may not attract a high enough market price.

The changes in abundance, size, and composition of shark and ray catch were attributed to a number of factors including overfishing and illegal fishing practices, among others. These factors, which fishers perceived as causing changes in shark and ray stock status, reflected the threats faced by sharks in Kenya (Temple *et al.,* 2019) and the UAE (Jabado, 2014). Globally, overfishing is recognized as the prominent threat to shark and ray species (Dulvy *et al.,* 2014). The biological attributes of sharks, which generally include slow growth rates, late sexual maturity, low reproductive rates and long life spans make them exceptionally vulnerable to overfishing and population decline (Myers and Ottensmeyer, 2005; Klimley, 2013). Overfishing has already been reported to be the major threat to marine fishery resources in Ghana (Overå, 2011). In Ghana, overfishing is influenced mainly through over-capitalization of the fishing industry, use of small mesh nets, human population pressure, increasing demand for fish products, and open-access regimes (Nunoo *et al.,* 2014). The historical increase in the number of both artisanal and industrial fleets in the early 2000s further worsened the depletion of Ghanaian fisheries resources (Lam *et al.,* 2012). Within these periods, the number of industrial fleets in Ghana nearly doubled, while artisanal canoes increased to about 24% (Anon, 2003). Many fishers were also concerned about the practices of illegal, unreported and unregulated (IUU) fishing, specifically stating light fishing and dynamiting as the major cause of decline in their target species. Despite the increasing efforts by the national Fisheries Commission and conservation organizations in educating fishers on the demerit of these methods of fishing, most fishers still believe the only available remedy to get abundant catch is to use these methods (Afoakwah *et al.,* 2018). Illegal fishing violates management efforts and is recognized as a serious threat to the sustainability of fisheries due to the adverse impact on the ecology of the oceans and economy of fishing nations (Okafor-Yarwood, 2019). Further, some fishers attributed much of the illegal, unreported and unregulated fishing practices in the region to the actions of foreign vessels fishing in the exclusive economic zone of coastal zones in Ghana, while most fishers also recognized illegal, unreported and unregulated fishing as common practices in Ghanaian fishing vessels.

## 5. Conclusions and recommendations

The study revealed that six main ecological variables shape fishing operations of artisanal fishers in Ghana. These comprise the season and weather conditions, lunar phase, bait type, seabird presence and fish movement, color of seawater, and sea current. Second, fishers have noticed a substantial decrease in shark and ray abundance and attributed the decline in numbers and size, as well as the species composition of their catch to overfishing and illegal, unreported and unregulated fishing. Finally, fishers noticed the depletion of some species as early as the 2000s, while most sharks and rays were observed to be depleted by the 2010s. The LEK of the studied fishers were consistent with empirical scientific studies. These baseline data are particularly essential in the design and implementation of policies and programs aimed at making elasmobranch fisheries economically and environmentally sustainable. Our data are beneficial to data poor fisheries research and management in Ghana and other countries that have limited funds to conduct fishery research that rely on traditional scientific surveys.

Our results show that the abundance of numerous shark and ray species has significantly depleted and fishers are aware of changes in the stock status. This calls for active management to mitigate this impact on sharks. Such management may require changes in and enforcement of gear regulations and effective monitoring of practices and activities of fishers, especially checking illegal, unreported and unregulated practices in both artisanal and foreign fleets. We further recommend the collaboration between research scientists, policymakers and artisanal fishers in any research, decision making, and management interventions as the latter possess much useful information of which the former may not be aware.

## Acknowledgements

Special thanks go to the Zoological Society of London EDGE of Existence Program team members for their training, advice and mentoring. My heartfelt appreciation to Benard Seret for his immense contribution, mentoring, advice and guidance for the species identifications and towards the successful completion of the study. Finally, the Authors appreciation goes to Moro Seidu, Bukari Saphianu, Paul Tehoda, Emmanuel Amoah, Clement Sullibie Saagulo Naabeh and Adomako Ohene as well as the local volunteers Isaac Assefuah, Kingford Amankwah, Jona Aquah, Timothy Amiah and Ben Adjei for their role in field data gathering. We are grateful to the staff of the Western Regional Fisheries Commission, chiefs, Chief fishers and their elders, fishers, traders and people of the five study communities for their cooperation and support in making this study possible.

## References

Afoakwah, R., Osei, M. B. D., and Effah, E. 2018. A Guide on Illegal Fishing Activities in Ghana. USAID/Ghana Sustainable Fisheries Management Project. Narragansett, RI: Coastal Resources Center, Graduate School of Oceanography, University of Rhode Island. Prepared by the University of Cape Coast, Ghana. GH2014_SCI048_UCC.

Aheto, D. W., Asare, N. K., Quaynor, B., Tenkorang, E. Y., Asare, C., Okyere, I. 2012. Profitability of small-scale fisheries in Elmina, Ghana. Sustainability, 4(11), 2785–2794.

Ainsworth, C. H., Pitcher, T. J., and Rotinsulu, C. 2008. Evidence of fishery depletions and shifting cognitive baselines in Eastern Indonesia. Biological Conservation, 141(3), 848–859.

Alexander, L., Agyekumhene, A., and Allman, P. 2017. The role of taboos in the protection and recovery of sea turtles. Frontiers in Marine Science, 4, 237.

Amador, K., Bannerman, P., Quartey, R., and Ashong, R. 2006. Ghana canoe frame survey 2004. Marine Fisheries Research Division, Accra, Technical Paper (34), 43.

Anadón, J. D., Giménez, A., Ballestar, R., and Pérez, I. 2009. Evaluation of local ecological knowledge as a method for collecting extensive data on animal abundance. Conservation biology, 23(3), 617–625.

Anon. 2003. China Customs Statistics Yearbook (1998–2000 data). General Administration of Customs of the People’s Republic of China. FIGIS Country Profile Factsheet. FAO, Rome.

Ballouard, J. M, Robillard, M., and Yvon, C. 2006. Statut et conservation des poissons-scies et autres chondrichtyens menacés en Afrique de l’Ouest: Rapport de synthèse. Rapport Missions 2005 et 2006, Poissons-scies. Noé Conservation – CSRP.

Berkes, F., Colding, J., and Folke, C. (Eds.). 2008. Navigating social-ecological systems: building resilience for complexity and change. Cambridge University Press.

Bonfil, R. 1994. Overview of world elasmobranch fisheries (No. 341). Food and Agriculture Organization.

Bräutigam, A., Callow, M., Campbell, I. R., Camhi, M. D., Cornish, A. S., Dulvy, N. K., Fordham, S. V., Fowler, S. L., Hood, A. R., McClennen, C., Reuter, E. L., Sant, G., Simpfendorfer, C. A., and Welch, D. J. 2015. Global Priorities for Conserving Sharks and Rays: A 2015–2025 Strategy.

Buckley, L., and Hile, J. 2007. End of the Line? (Second Edition). WildAid and Oceana. http://oceana.org/sites/default/files/o/fileadmin/oceana/uploads/Sharks/EndoftheLine_Spread_sm.pdf

Castillo-Géniz, J. L., Márquez-Farias, J. F., De La Cruz, M. R., Cortés, E., and Del Prado, A. C. 1998. The Mexican artisanal shark fishery in the Gulf of Mexico: towards a regulated fishery. Marine and Freshwater Research, 49(7), 611–620.

Clarke, S. 2005. Trade in shark products in Singapore, Malaysia and Thailand. Singapore, Southeast Asian Development Center and ASEAN.

Cook, S. F. 1990. Trends in Shark Fin Markets: 1980, 1990, and Beyond. Chondros, 3–6.

CRC. 2013. Solving the Fisheries Crisis in Ghana: A Fresh Approach to Collaborative Fisheries Management, USAID-URI Integrated Coastal and Fisheries Governance (ICFG) Initiative. Coastal Resources Center, Graduate School of Oceanography, University of Rhode Island. 20p.

deGraft-Johnson, K. A. A., Blay, J., Nunoo, F. K. E., and Amankwah, C. C. 2010. Biodiversity threats assessment of the Western Region of Ghana. The integrated coastal and fisheries governance (ICFG) initiative Ghana.

Demuynck, K. 1994. The participatory rapid appraisal on perceptions and practices of fisherfolk on fishery resource management in an artisanal fishing community in Cameroon. Food and Agriculture Organization.

Dulvy, N. K., and Polunin, N. V. C. 2004. Using informal knowledge to infer human-induced rarity of a conspicuous reef fish. Animal Conservation 7, 365–374.

Dulvy, N. K., Fowler, S. L., Musick, J. A., Cavanagh, R. D., Kyne, P. M., Harrison, L. R.,…and Pollock, C. M. 2014. Extinction risk and conservation of the world’s sharks and rays. elife, 3, e00590.

Dulvy, N. K., Simpfendorfer, C. A., Davidson, L. N. K., Fordham, S. V., Bräutigam, A., Sant, G. and Welch, D. J. 2017. Challenges and Priorities in Shark and Ray Conservation. Current Biology 27, R565–R572.

Eddy, T. D., Gardner, J. P., and Pérez-Matus, A. 2010. Applying fishers’ ecological knowledge to construct past and future lobster stocks in the Juan Fernández Archipelago, Chile. PLoS One, 5(11), e13670.

Fabinyi, M. 2012. Historical, cultural and social perspectives on luxury seafood consumption in China. Environmental Conservation, 39(1), 83–92.

FAO. 2010. Report of the FAO FishCode-STF/CECAF/FCWC Subregional Workshop on the Improvement of Fishery Information and Data Collection Systems in the West Central Gulf of Guinea Region. Accra, Ghana, 26-28 June 2007. Available online: www.fao.org/docrep/012/k7479b/k7479b00.pdf

Finegold, C., Gordon, A., Mills, D., Curtis, L., and Pulis, A. 2010. Western region fisheries sector review. World Fish Center. USAID Integrated Coastal and Fisheries Governance Initiative for the Western Region, Ghana.

Gelber, M. J. 2018. Plenty of Fish in the Sea? Shark Fishing and the Fin Trade in Ghana: A Biting Review. Master Degree Thesis. University of Florida, in Gainesville, FL USA.

Ghana Statistical Services. 2014. 2010 Population and housing census: District analytical report. Ghana: Accessed June, 8, 2019.

Gomes, C., Mahon, R., Hunte, W., and Singh-Renton, S. 1998. The role of drifting objects in pelagic fisheries in the Southeastern Caribbean. Fisheries Research, 34(1), 47–58.

Grant, S., and Berkes, F. 2007. Fisher knowledge as expert system: A case from the longline fishery of Grenada, the Eastern Caribbean. Fisheries Research, 84(2), 162–170.

Jabado, R. W. 2014. Assessing the fishery and ecology of sharks in the United Arab Emirates. PhD Dissertation. United Arab Emirates University. UAE.

Jaiteh, V. F., Hordyk, A. R., Braccini, M., Warren, C., and Loneragan, N. R. 2017. Shark finning in eastern Indonesia: assessing the sustainability of a data-poor fishery. ICES Journal of Marine Science, 74(1), 242–253.

Johannes, R. E., and Neis, B. 2007. The value of anecdote. Fishers’ Knowledge in Fisheries Science and Management. Haggan, N., Neis, B. and Baird, IG (eds). UNESCO, Paris, 41–58.

Johannes, R. E., Freeman, M. M., and Hamilton, R. J. 2000. Ignore fishers’ knowledge and miss the boat. Fish and Fisheries, 1(3), 257–271.

Kalikoski, D. C., and Vasconcellos, M. 2007. The role of fishers’ knowledge in the co-management of small-scale fisheries in the estuary of Patos Lagoon, Southern Brazil. Fishers’ Knowledge in fisheries science and management, 289–312.

Kelleher, K. 2005. Discards in the world’s marine fisheries: an update. FAO Fisheries Technical Paper 470. Food and Agriculture Organization (FAO), Rome, Italy.

Klimley, A. P. 2013. The biology of sharks and rays. University of Chicago Press.

Kyne, P. M., Jabado, R. W., Rigby, C. L., Dharmadi, Gore, M. A., Pollock, C. M., Herman, K. B., Cheok, J., Ebert, D. A., Simpfendorfer, C. A., and Dulvy, N. K. 2020. The thin edge of the wedge: extremely high extinction risk in wedgefishes and giant guitarfishes. Aquatic Conservation - Marine and Freshwater Ecosystems, 1–25.

Lam, V. W., Cheung, W. W., Swartz, W., and Sumaila, U. R. 2012. Climate change impacts on fisheries in West Africa: implications for economic, food and nutritional security. African Journal of Marine Science, 34(1), 103–117.

Lam, V. Y. Y., and Sadovy de Mitcheson, Y. 2011. The sharks of South East Asia – unknown, unmonitored and unmanaged. Fish and Fisheries, 12(1), 51–74.

Lunn, K. E., and Dearden, P. 2006. Monitoring small-scale marine fisheries: An example from Thailand’s Ko Chang archipelago. Fisheries Research, 77(1), 60–71.

Mangel, M., Talbot, L. M., Meffe, G. K., Agardy, T. M., Alverson, D. L., Barlow, J., et al. 1996. Principles for the conservation of wild living resources. Ecological Applications, 6(2), 338–362.

Manire, C. A., and Gruber, S. 1990. Many sharks may be headed toward extinction. Conservation Biology, 4, 10–11.

McCleery, R. A., Ditton, R. B., Sell, J., and Lopez, R. R. 2006. Understanding and improving attitudinal research in Wildlife Sciences. Wildlife Society Bulletin, 34(2), 537–541.

McCool, S. F., and Guthrie, K. 2001. Mapping the dimensions of successful public participation in messy natural resources management situations. Society and Natural Resources, 14(4), 309 – 323.

Miller, K. K. 2009. Human dimensions of wildlife population management in Australasia - history, approaches and directions. Wildlife Research, 36, 48–56.

Ministry of Fisheries and Aquaculture Development 2015. National Fisheries Management Plan, Government of Ghana pp 48.

Moore, J. E., Cox, T. M., Lewison, R. L., Read, A. J., Bjorkland, R., McDonald, S. L.,…and Joynson-Hicks, C. 2010. An interview-based approach to assess marine mammal and sea turtle captures in artisanal fisheries. Biological Conservation, 143(3), 795–805.

Myers, R. A., and Ottensmeyer, C. A. 2005. Extinction risk in marine species. In ‘Marine Conservation Biology: the Science of Maintaining the Sea’s Biodiversity’.(Eds EA Norse and LB Crowder.) pp. 126–174.

Naderifar, M., Goli, H., and Ghaljaie, F. 2017. Snowball sampling: A purposeful method of sampling in qualitative research. Strides in Development of Medical Education, 14(3), 1–6.

Newing, H. 2010. Conducting research in conservation: social science methods and practice. Routledge.

Nunoo, F. K. E., Asiedu, B., Amador, K., Belhabib, D., and Pauly, D. (2014). Reconstruction of marine fisheries catches for Ghana, 1950–2010. Vancouver (Canada): Fisheries Centre, University of British Columbia. http://www.seaaroundus.org/doc/publications/wp/2014/Nunoo-et-al-Ghana.pdf P. 2.

Okafor-Yarwood, I. 2019. Illegal, unreported and unregulated fishing, and the complexities of the sustainable development goals (SDGs) for countries in the Gulf of Guinea. Marine Policy, 99, 414–422.

Olsson, P., and Folke, C. 2001. Local ecological knowledge and institutional dynamics for ecosystem management: a study of Lake Racken watershed, Sweden. Ecosystems, 4(2), 85–104.

Overå, R. 2011. Modernisation narratives and small-scale fisheries in Ghana and Zambia. In Forum for Development Studies (Vol. 38, No. 3, pp. 321–343). Routledge.

Oxenford, H. A. 1999. Biology of the dolphinfish (Coryphaena hippurus) in the western central Atlantic: a review. Scientia Marina, 63(3-4), 277–301.

Poizat, G., and Baran, E. 1997. Fishermen’s knowledge as background information in tropical fish ecology: a quantitative comparison with fish sampling results. Environmental Biology of fishes, 50(4), 435–449.

Rasalato, E., Maginnity, V., and Brunnschweiler, J. M. 2010. Using local ecological knowledge to identify shark river habitats in Fiji (South Pacific). Environmental Conservation, 37(1), 90–97.

Riley, S. J., Decker, D. J., Carpenter, L. H., Organ, J. F., Siemer, W. F., Mattfeld, G. F., et al. 2002. The essence of wildlife management. Wildlife Society Bulletin, 30(2), 585–593.

Robillard, M. and Séret, B. 2006. Cultural importance and decline of sawfish (Pristidae) populations in West Africa. Cybium 30: 23–30.

Ruddle, K. 2000. Systems of knowledge: dialogue, relationships and process. Environment, development and Sustainability, 2: 277–304.

Salas, S., Chuenpagdee, R., Seijo, J. C., and Charles, A. 2007. Challenges in the assessment and management of small-scale fisheries in Latin America and the Caribbean. Fisheries research, 87(1), 5–16.

Silva, G. 2005. The classification of living beings among the fishermen of Piratininga, Rio de Janeiro dlm. Diugues, A.C. (pytg.) Maritime Anthropology in Brazil. Sao Paulo (76–79).

Silvano, R.A.M. and Valbo-Jorgensen, J. 2008. Beyond fishermen’s tales: contribution of fishers’ local ecological knowledge to fish ecology and fisheries management. Environment, Development and Sustainability, DOI 10.1007/s10668-008-9149-0.

Silver, J. J., and Campbell, L. M. 2005. Fisher participation in research: dilemmas with the use of fisher knowledge. Ocean and Coastal Management, 48(9-10), 721–741.

Taylor, R. B., Morrison, M. A., and Shears, N. T. 2011. Establishing baselines for recovery in a marine reserve (Poor Knights Islands, New Zealand) using local ecological knowledge. Biological Conservation, 144(12), 3038–3046.

Temple, A. J., Wambiji, N., Poonian, C. N., Jiddawi, N., Stead, S. M., Kiszka, J. J., and Berggren, P. 2019. Marine megafauna catch in southwestern Indian Ocean small-scale fisheries from landings data. Biological conservation, 230, 113–121.

Vlietstra, L. S. 2005. Spatial associations between seabirds and prey: effects of large-scale prey abundance on small-scale seabird distribution. Marine Ecology Progress Series, 291, 275–287.

Wing, S. R. and Wing, E. S. 2001. Prehistoric fisheries in the Caribbean. Coral Reefs, 20, 1–8.

